# Quantitative single cell heterogeneity profiling of patient derived tumor initiating gliomaspheres reveals unique signatures of drug response and malignancy

**DOI:** 10.1101/2020.01.14.900506

**Authors:** Michael Masterman-Smith, Nicholas A. Graham, Ed Panosyan, Jack Mottahedeh, Eric E. Samuels, Araceli Nunez, Sung Hyun Lim, Tiffany Phillips, Meeryo Choe, Koppany Visnyei, William H. Yong, Thomas G. Graeber, Ming-Fei Lang, Harley I. Kornblum, Jing Sun

**Author notes:** Corresponding author: Jing Sun. Additional File #1: Raw Dataset. Additional File#2: Supplementary Methods.

## Abstract

**Background:** Glioblastoma is a deadly brain tumor with median patient survival of 14.6 months. At the core of this malignancy are rare, highly heterogenous malignant stem-like tumor initiating cells. Aberrant signaling across the EGFR-PTEN-AKT-mTOR signal transduction pathways are common oncogenic drivers in these cells. Though gene-level clustering has determined the importance of the EGFR signaling pathway as a treatment indicator, multiparameter protein-level analyses are necessary to discern functional attributes of signal propagation. Multiparameter single cell analyses is emerging as particularly useful in identifying such attributes.

**Methods:** Single cell targeted proteomic analysis of EGFR-PTEN-AKT-mTOR proteins profiled heterogeneity in a panel of fifteen patient derived gliomaspheres. A microfluidic cell array ‘chip’ tool served as a low cost methodology to derive high quality quantitative single cell analytical outputs. Chip design specifications produced extremely high signal-to-noise ratios and brought experimental efficiencies of cell control and minimal cell use to accommodate experimentation with these rare and often slow-growing cell populations. Quantitative imaging software generated datasets to observe similarities and differences within and between cells and patients. Bioinformatic self-organizing maps (SOMs) and hierarchical clustering stratified patients into malignancy and responder groups which were validated by phenotypic and statistical analyses.

**Results:** Fifteen patient dissociated gliomaspheres produced 59,464 data points from 14,866 cells. Forty-nine molecularly defined signaling phenotypes were identified across samples. Bioinformatics resolved two clusters diverging on EGFR expression (*p* = 0.0003) and AKT/TORC1 activation (*p* = 0.08 and *p* = 0.09 respectively). TCGA status of a subset showed genetic heterogeneity with proneural, classical and mesenchymal subtypes represented in both clusters. Phenotypic validation measures indicated drug responsive phenotypes to EGFR blocking were found in the EGFR expressing cluster. EGFR expression in the subset of drug-treated lines was statistically significant (p<.05). The EGFR expressing cluster was of lower tumor initiating potential in comparison to the AKT/TORC1 activated cluster. Though not statistically significant, EGFR expression trended with improved patient prognosis while AKT/TORC1 activated samples trended with poorer outcomes.

**Conclusions:** Quantitative single cell heterogeneity profiling resolves signaling diversity into meaningful non-obvious phenotypic groups suggesting EGFR is decoupled from AKT/TORC1 signalling while identifying potentially valuable targets for personalized therapeutic approaches for deadly tumor-initiating cell populations.

## Introduction

The cell of origin for many cancers is a specific, rare subset of cancer cells responsible for tumor initiation, cellular heterogeneity and various features that underlie the malignant nature of the cancer types which they have been identified in. (1,2) Among those cancers with a cancer stem cell origin is the highly malignant and deadly glioblastoma brain tumor. Patients have a median survival of 14.6 months from diagnosis and five year survival is an abysmal 5%. (3)

Patient glioblastoma tumors yield relatively stable cancer stem-like, tumor initiating cell populations which retain some of the phenotypical and genetic heterogeneity of the cancers they produce *in vitro*. (4) Being able to recover these cells from patient biopsies *in vitro* are a robust predictor of clinical progression and outcome. These cells additionally serve as useful substrates for drug discovery and to determine the essential molecular signaling landscape contributing to *in vivo* malignancy and resistance. (5,6)

The mechanisms underlying gliomasphere malignancy can be defined by pathway redundancies in the biological systems controlling states of oncogenic activation in cancer cells and their cancer initiating subtypes. (4,5) Signaling along the EGFR-PTEN-AKT-TORC1 signaling axes provide phenotypic features in gliomasphere cell populations. (7) These pathways are especially important in governing cellular fate decisions by transmitting signals controlling survival, self-renewal, growth, proliferation, metabolism, glycolytic adaptation, drug efflux and symmetric division, among other essential features. (8) (9) (10) (11) Targets of EGFR signaling have long been the therapeutic and diagnostic targets of glioblastoma which extends into the era of cancer immunotherapy, which utilizes EGFR in chimeric antigen receptor (CAR) T-cell therapy. (12,13)

Clustering gliomasphere models according to The Cancer Genome Atlas (TCGA) classification system has provided insight into the genetic landscape of gliomaspheres. (10) Mesenchymal and non-mesenchymal gliomasphere subtypes have been delineated with non-mesenchymal gliomaspheres consisting of both classical and proneural subtypes. (11) Other studies have found gliomasphere models cluster according to a lower malignancy proneural and higher malignancy mesenchymal classification with proneural status conferring phenotypes with lower sphere formation and improved survival in *in vivo* xenografts of gliomaspheres. (14,15)

Based on the gene-level mutations, EGFR mutations (including point mutations, amplifications, rearrangements, and alternative splicing) are found in all subtypes of glioblastoma and are present in 57% of glioblastoma (7,16,17). At the protein level, cellular EGFR expression is tightly controlled in normal but not in cancerous cells by epigenetic regulation and protein degradation pathways, leading to overall high EGFR protein levels (18,19). These findings indicate that the gene-level mutations of EGFR and its protein-level expression can be vastly different.

Given the discrepancies between the gene-level mutations and the protein-level expression of EGFR and other molecules in the EGFR signaling pathways, it is important to detect the protein level changes in gliomasphere. Multiparameter single cell measurement of EGFR, PTEN, activated AKT and TORC1 signaling had been used previously to aggregate and identify prognostic glioblastoma subtypes. (20) By extending this methodology across a panel of patient-derived gliomasphere samples, we sought to observe and detail the signaling diversity within this stem-like subset of cells. In measuring this native signaling heterogeneity and deploying cluster-based analyses with comparison to genotypic and phenotypic descriptors, features of response and target characterization can be observed.

## Results

### Experimental design

Patient glioblastoma tumors **(Fig. 1A**, **left)** were dissociated and placed in defined serum free enrichment media to select for *in vitro* growth and expansion of gliomaspheres as neurospheres **(Fig. 1A**, **middle, right)**. After stem-like cell selection and enrichment, neurospheres are dissociated into single cells and loaded into chambers of microfluidic cell array chips for quantitative immunocytochemical staining and imaging **(Fig. 1B**, **left, Supplementary Methods).** Imaging software quantifies the average fluorescent intensity from each cell for each defined biomarker as the means to reflect individual cellular protein concentration **(Fig. 1B**, **middle)**. Bioinformatic analysis of a dataset of all cells from a series of samples resolves complex intra- and -inter sample signaling heterogeneity. The resulting data is validated with genotypic and phenotypic measures to assess functional status (**Fig. 1C**, **bottom right)**.

**Figure 1.**
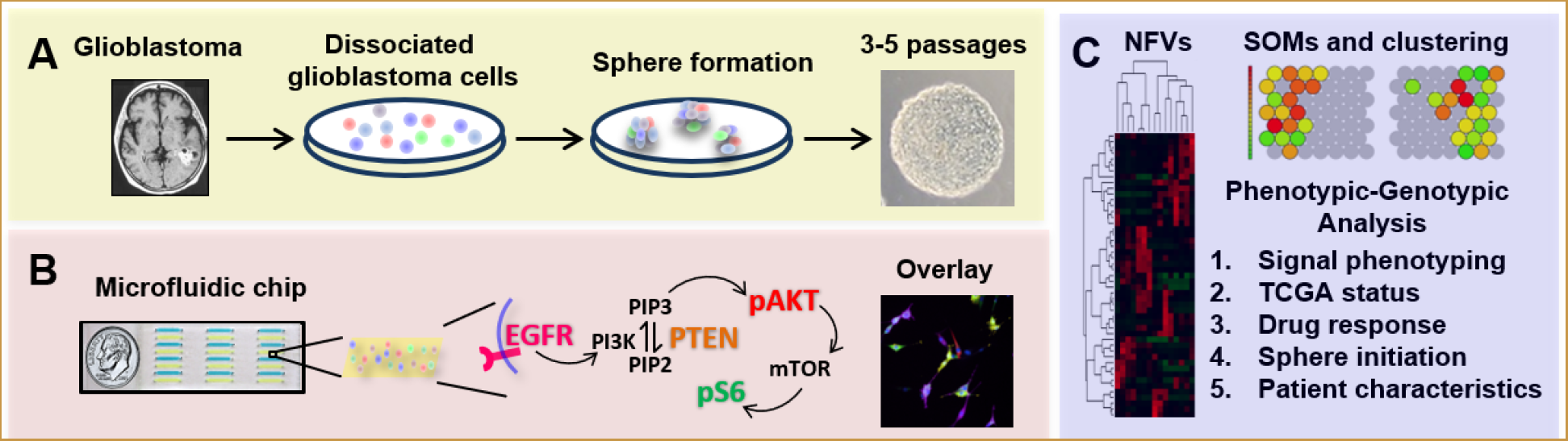
Conceptual summary of quantitative single cell heterogeneity profiling of patient-derived glioblastoma tumor initiating cells (TICs). A. (Left) T1 weighted MRI of brain tumor in left parieto-occipital lobe (in white). (Middle) Clinical glioblastoma samples are dissociated into single cells and placed in ‘neurosphere’ serum-free enrichment media for in vitro selection and expansion of TICs. (Right) Brightfield image of classic 3D gliomasphere cell line sphere. Colored cells illustrate cellular heterogeneity. B. (Left) Image of 24 chamber microfluidic device. Chambers are filled with alternating yellow and blue dyes for visualization. Glioblastoma TIC are dissociated into single cells, loaded into microfluidic channels, labelled with fluorophore-conjugated antibodies for four signaling proteins (EGFR, PTEN, pAKT, and pS6) to measure signal through the oncogenic EGFR/PI3K/AKT/mTOR pathways. [anti-EGFR (purple), anti-PTEN (orange), anti-pAKT (red), and anti-pS6 (green)]. (Right) Fluorescent signal intensity of each the four markers of each cell is via quantification of immunofluorescent signal intensities of labelled single cells. C. Quantitative analysis involves: (Left) Unsupervised hierarchical clustering of neighborhood frequency vectors (NFVs) derived from ~1000 cells of each sample’s individual self-organizing map (SOM). (Top, right) Resultant self-organizing maps (SOMs) of the two clusters of gliomaspheres. (Lower, right) Phenotypic-genotypic analysis of clustering included validation by: signal phenotyping, The Cancer Genome Atlas (TCGA) subgrouping, sphere-based drug response measures, sphere initiation efficiency, and patient characteristics of patient survival and disease progression.

### Heterogeneity profiling of patient-derived gliomaspheres

A series of bioinformatic steps quantified individual and multiparameter, parallel datasets to characterize and discern cellular heterogeneity among cells and patients.․ In total, 14,866 cells (mean = 991 cells/line), produced 59,464 individual data points from fifteen human gliomasphere lines **(Supplemental Table 1)**. Pairwise average linkage using Pearson correlation clustering of individual protein expression vectors EGFR, PTEN, pAKT and pS6 identified reasonable correlation coefficients between PTEN and pS6 and pAKT and pS6 **(Fig. 2A)**. Boxplots of single cell expression of these markers for each sample revealed unique sample diversity and substantial molecular and cellular heterogeneity (Fig. 2B). Boxplots of all cells showed the spread of values for each marker in the dataset, showing, while individual parameters were not skewed, there existed a wide distribution of values for each marker **(Fig. 2B).** Self-organizing maps (SOMs) resolved forty-nine unique molecular phenotypes across patients **(Fig. 2C)**. Unsupervised hierarchical clustering based on neighborhood frequency vectors (NFVs) of self-organizing map (SOM) projections in Fig. 2C yielded two predominant clusters **(Fig. 3B-3C)**. By taking the average biomarker intensity of all cells in each cluster, two quantitative multiparameter signaling phenotypes emerged **(Fig. 3C**, **Supplemental Methods)**.

**Figure 2.**
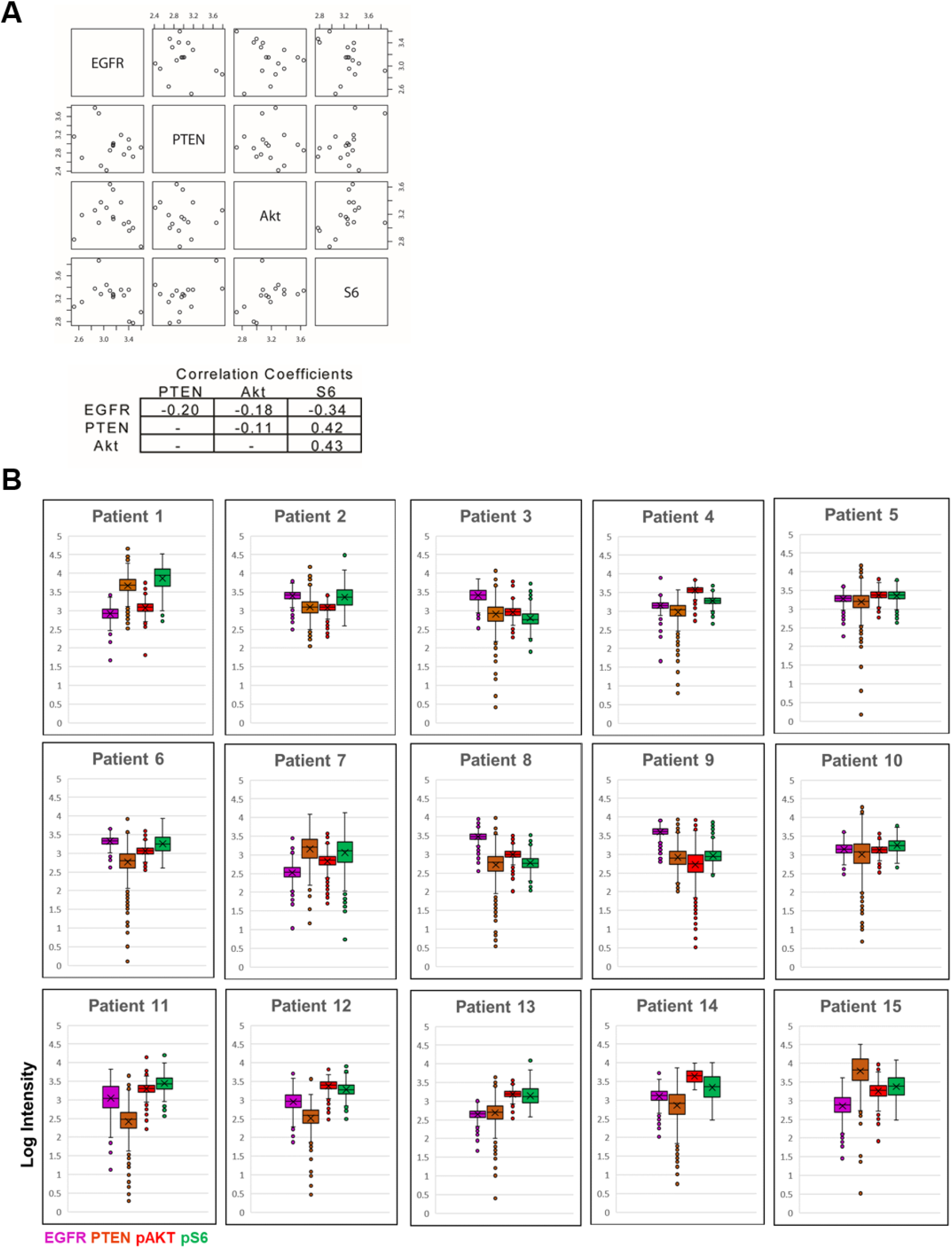
Single cell measurements of EGFR, PTEN, pAKT and pS6 across patients. **A.** Pairwise average linkage using the Pearson correlation clustering of individual mean protein expression vectors EGFR, PTEN, pAKT and pS6 show correlation coefficients between PTEN and pS6 and pAKTand pS6. **B.** Boxplot distribution of the complete dataset for each marker of all cells. Minimum non-outlier value is bottom horizontal line, first quartile represents bottom box, median is middle horizontal line denoted by an ‘X’, third quartile represents top box, and maximum non-outlier value is top horizontal line. Circles represent outlier values. Left, EGFR, pink. Middle left, PTEN, orange. Middle right, pAKT, red. Right, pS6, green.

**Figure 3.**
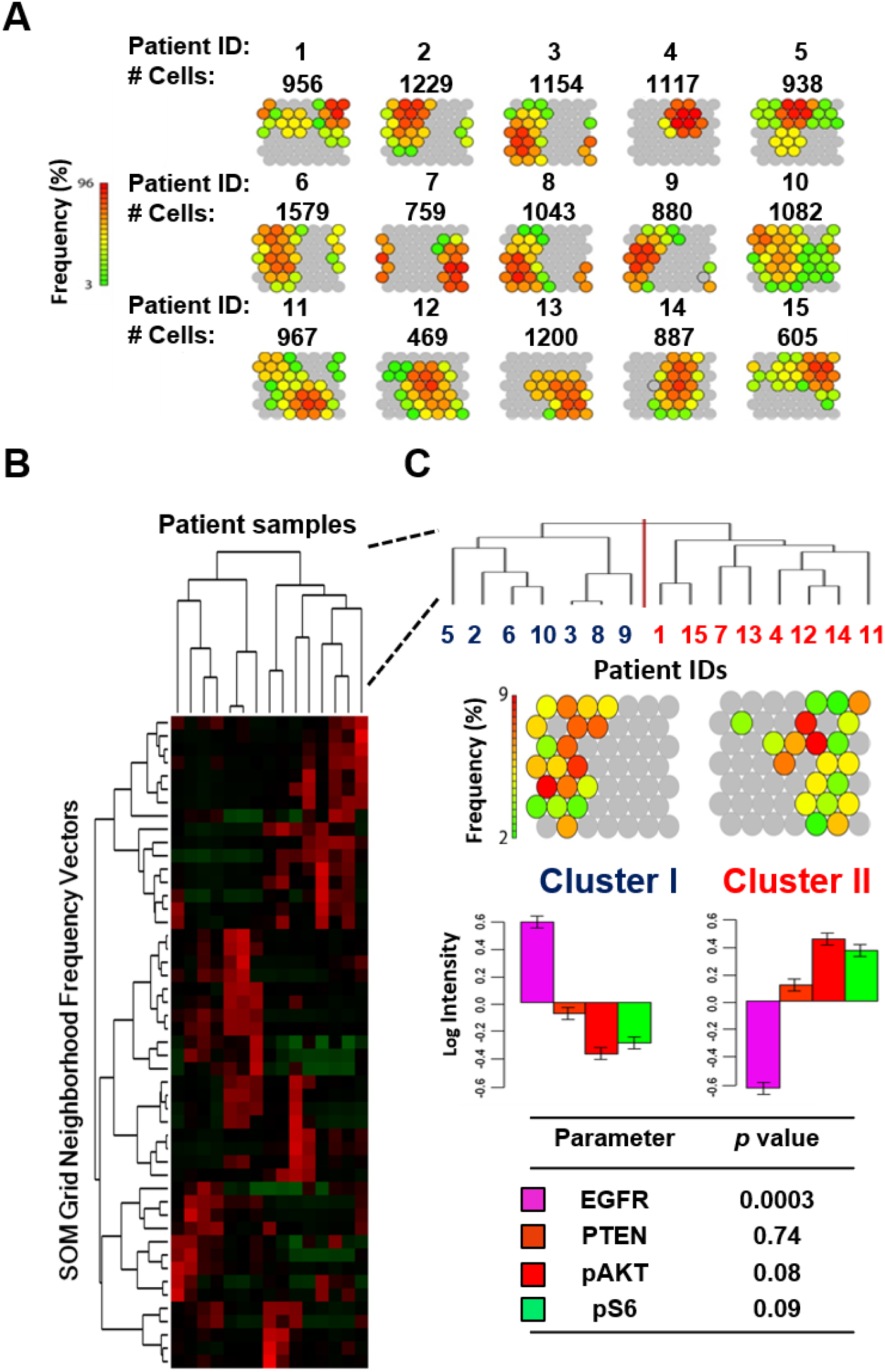
Multiparameter self-organizing map (SOM) and clustering of human derived gliomaspheres. A. Rainbow colored normalized self organizing maps (SOM) display groupings of forty-nine molecular phenotypes on a 7 × 7 grid for each of the 15 patients. Red, high percentage of cells. Green, low percentage of cells. Grey, <2% of cells. Above each patient map is the number of individual cells analyzed. Total cells analyzed=14,866. Range 469-1579 cells. Mean=991 cells. **B.** SOM derived Neighborhood Frequency Vectors (NFVs) for each of the 15 human-derived brain gliomasphere lines after unsupervised hierarchical clustering. A heatmap of the dataset where each row corresponds to one of 49 SOM unit and each column represents a patient gliomasphere line. Red and green indicate relative high and low neighborhood frequencies, respectively. **C.** (Top) Enlarged dendrogram reveals two main clusters. (Upper middle) Representative SOMs of each cluster. Color representation of the frequency at which individual cells are assigned to each SOM unit. (Lower middle) Waterfall plots of mean expression for each of the four markers in each cluster of PI3K/AKT/TORC1 pathways mapped. (Bottom) Student’s t-test on mean expression levels reveal significantly differentially expression of EGFR biomarker between each cluster with borderline significance for pAKT and pS6. EGFR, pink. PTEN, orange. pAKT, red. pS6, green.

**Figure 4.**
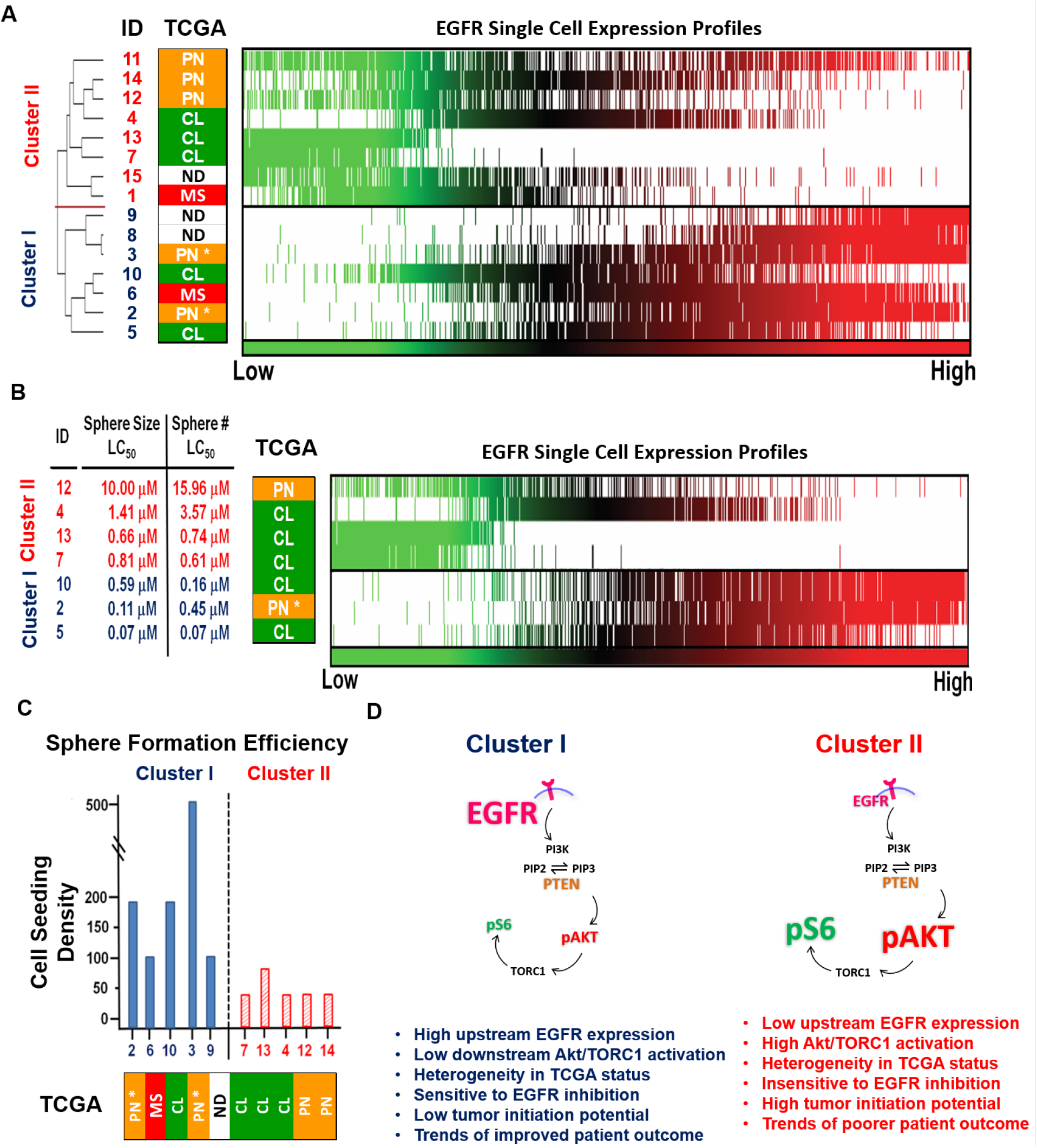
TCGA status, EGFR single cell profiling, drug response, malignancy profiles and activation schematic of clustered gliomasphere samples. **A.** (Left) The Cancer Genome Atlas (TCGA) groupings. Green, CL=Classical. Yellow, PN=Proneural. Red, MS=Mesenchymal. *IDH1 mutant. ND=No Data. (Right) Diversity of EGFR expression is observed by visualization heatmap of single cell profiling of EGFR expression for all 15 glioma cancer stem cell (gCSC) lines. Each vertical line corresponds to 1 cell set against a white background. Green=low expression, Red=high expression. **B.** LC_50_s based on sphere size and sphere number to EGFR inhibitor erlotinib from a random sampling of 3 gliomasphere lines from Cluster I and 4 gliomasphere lines from Cluster II. (Middle) TCGA groupings of treated samples. (Right) Visualization heatmap of single cell EGFR expression of treated samples. **C.** *In vitro* sphere forming efficiency in subsample of 4 gliomasphere lines from each cluster. Y-axis, number of cells seeded to form 10 neurospheres. (Mann Whitney U Test, two-tailed, *p* < 0.016, SD = 139.04). **D.** Schematic representation of two gliomasphere clusters. Signal phenotyping denoted a high EGFR expressing cluster (Cluster I, blue) and activated AKT/TORC1 cluster (Cluster II, red). Genomic analysis identified evidence of genetic heterogeneity in each cluster. LC50s derived from sphere size and sphere number with the EGFR inhibitor erlotinib revealed the EGFR expressing Cluster I to be a drug responsive phenotype. Lower sphere initiation efficiency and trends of improved outcome were observed in EGFR expressing Cluster I in comparison to activated AKT/TORC1 Cluster II. EGFR (purple), PTEN (orange), (red), and pS6 (green). Text size is indicator of level of expression.

Identified clusters revealed Cluster I to becharacterized by significantly high EGFR expression (*p* = 0.0003) with decreased pAKT and TORC1 in comparison to Cluster II, which had lower EGFR expression and higher pAKT and pS6 levels (*p* = 0.08, and *p* = 0.09 respectively). PTEN expression was statistically insignificant and barely discernible between clusters.

### TCGA grouping of gliomaspheres reveal genotypic heterogeneity in clusters

The Cancer Genome Atlas (TCGA) subgroupings were available on twelve of fifteen gliomasphere samples across the EGFR expressing Cluster I (5/7 samples, 71.4%) and AKTAKT/TORC1 activated Cluster II (7/8 samples, 87.5%) **(Fig. 3A)**. The EGFR expressing Cluster I had two proneural samples (2/5, 40.0%), two classical samples (2/5, 40.0%) and one mesenchymal sample (1/5, 20.0%). Isocitrate dehydrogenase 1 gene mutations (IDHR1324) were found in both proneural samples in this cluster (patients 2 and 3). (21) Within the AKTAKT/TORC1 activated Cluster II, there were three classical samples (3/7, 42.8%), three proneural samples (3/7, 42.8%) and one mesenchymal sample (1/7, 14.2%) **(Supplemental Table 2)**.

### High EGFR expressing Cluster I samples are responsive to EGFR inhibition

Visualization of single cell EGFR expression profiling revealed a broad diversity in EGFR expression **(Fig. 3A)**. A randomly selected subsample of seven gliomasphere lines, all of which were either proneural or classical samples, were tested for response to the EGFR blocker erlotinib **(Fig. 3B)**. LC_50_ measurements of sphere size and sphere number showed EGFR expression Cluster I had lower LC_50_ and high AKTAKT/TORC1 Cluster II had higher LC_50_s (Sphere size: Cluster I LC_50_ mean = 0.26μM, Cluster II LC_50_ mean = 3.27μM. Sphere number: Cluster I LC_50_ mean = 0.33μM, Cluster II LC_50_ mean = 5.22μM). Non-parametric Mann-Whitney U tests on LC_50_s of sphere size and sphere number showed significant response to EGFR blockade in EGFR expressing Cluster I in comparison to AKTAKT/TORC1 activated Cluster II (Sphere size, *p* = 0.029, SD = 3.57, two-tailed; Sphere number *p* = 0.029, SD = 5.81). Though significant differences in sphere size and sphere number were found, the large standard deviation prohibits definitive discrimination between clusters. Significant differences were found in mean EGFR expression and borderline significance in median EGFR expression within this subsample of drug-treated lines, indicating a relationship between receptor expression and drug response (*p* =0.0418 mean, 0.0501 median, T test, unpaired, 2-sided, unequal variance) **(Supplemental Information)**.

### Molecularly defined clusters differed in malignancy response

Sphere formation efficiency was measured as an *in vitro* means to assess tumor initiation potential in a random subsample of five gliomasphere lines from each cluster which included proneural, classical and mesenchymal samples. **(Fig. 3C)**. Non-parametric Mann-Whitney U tests of sphere formation efficiency showed significantly higher sphere formation efficiency in AKTAKT/TORC1 activated Cluster II in comparison to EGFR expressing Cluster I (*p* =.0159, SD = 139.04). Though statistically significant, the standard deviation was quite large for definitive discrimination between clusters.

Kaplan-Meier curves were generated based on progression free survival and overall survival of patients gliomasphere lines were derived from **(Supplementary Figure 4b).** Though there were trends of better prognosis in EGFR expressing Cluster I and poorer prognosis in AKT/TORC1 activated Cluster II, hazard ratios for these outcome measures were not statistically significant in predicting prognosis.

## Discussion

Single cell analysis for the purposes of cellularly heterogeneity profiling is becoming increasingly relevant for diagnostics, drug discovery, preclinical drug development, and basic and translational research. (20,22) It is an important methodology for the dissemination and categorization of complex cellular heterogeneity and to improve life sciences research and development in fields where rare cells may be involved in natural and disease processes. These sensitive analyses are becoming particularly relevant in cancer stem cell biology to understand the extreme cellular, molecular and genetic heterogeneity and additionally identify potential targetable cell populations. (23,24) Single cell datasets often achieve a greater resolution than uniparameter analysis because, embedded in the methodology, are detailed examinations of the relationships between nodes in the oncogenic networks studied. With the increasing incorporation of sensitive multiparameter single cell analysis technologies such as CyTOF and DNA barcoding to study putative tumor initiating cells, the methodologies have been deployed to observe putative brain cancer stem cells in parental tumors and characterize inherent molecular and functional heterogeneity and fates of these elusive yet highly malignant cells. (25,26)

Patient derived spheroid model systems may help define targetable cell subpopulations responsible for tumor initiation and malignancy. However, the cells, in the two decades since their identification, remain ambiguous to characterize and therapeutic interventions, for the range of malignant cancers they have been proposed to initiate, are still a challenge. In this study, we sought to focus on the contribution of EGFR signaling on downstream AKT-TORC1 signaling pathways in gliomaspheres to discern inter- and intra- sample similarities and differences in these pathways.

LC_50_s of EGFR inhibition based on sphere number and sphere size revealed EGFR expressing Cluster I samples to be drug responsive in comparison to AKTAKT/TORC1 activated Cluster II cells. Mean EGFR expression of treated samples differed significantly between clusters indicating a predictive target-response relationship of EGFR expression to inhibition.․ This finding is in itself important as the evolution of computational target-response modelling is becoming an important component in drug development to screen out potential candidate failures as early as possible and move efficacious candidates to market quicker at reduced costs. The finding is consistent with other EGFR blocking studies in patient derived gliomasphere models in achieving reduced sphere growth. It alsolends downstream mechanistic insight into why some gliomaspheres grow and proliferate in the absence of exogenous mitogen EGF supplementation and recover from EGFR blocking action. (27)

*In vitro* sphere formation assays, as a measure of tumor initiation potential, revealed the high EGFR expressing Cluster I to have lower sphere forming efficiency in comparison AKT/TORC1 activated signature. A Within the EGFR expressing samples a range of sphere formation efficiencies were found suggesting potential heterogeneity within this population. And although AKT/TORC1 had high sphere forming potential suggesting greater malignancy and perhaps complexity, the samples revealed uniform phenotypes on this measure.

Cluster based heterogeneity profiling did not significantly predict progression free survival or overall survival in patients **(Supplemental Figure 4b)**. Perhaps this may be due to a low number of samples or limited number of parameters. However, the high EGFR Cluster I showed trends of improved prognosis and AKT/TORC1 activated Cluster II trended towards poorer prognosis. Taken together with *in vitro* sphere efficiency assays, the methodology employed suggest these signaling phenotypes may play a role in meaningfully distinguishing populations. It has been found AKT and TORC1 activation are key drivers of malignancy, reactivation, treatment resistance and response phenomena. (28) Thus, the identity of a therapeutic foci in gliomaspheres may come to be of use in modeling interventions at these cells. (29)

The clusters consisted of proneural, classical and mesenchymal TCGA groupings. All EGFR expressing Cluster I proneural gliomaspheres had mutations in the isodehyrodgenase isocitrate dehydrogenase (IDH1) gene, while all AKT/TORC1 activated Cluster II proneural gliomaspheres did not harbor this mutation. Given IDH1 mutations are a feature of lower grade brain tumors, consistent with the sphere forming ability of these samples. Additionally, through recent evidence from other single cell analytical techniques, IDH1 mutation has been observed as a feature of EGFR amplification, suggesting this clustering is reasonable. (30)

It is relevant to recognize bulk, uniparameter or quantitatively insensitive cell metrics could have obscured subtle yet crucial, informative molecular differences in the cells which multiparameter single cell clustered datasets resolve. Correlative analysis of individual proteins, while identifying potentially meaningful correlations, could not reveal the level of cellular complexity this type of multiparameter, parallel analyses was able to discern in terms of the phenotypes identified. Of note, while some parameters studied did not reach statistical significance, they were still vital to resolving phenotype clusters. The activated states of AKT and TORC1, while of only borderline statistical significance, proved essential to distinguishing clusters. This is an important consideration given dual inhibition of these targets has shown evidence of therapeutic potential. This additionally supports an emerging understanding of low TORC1 activation is a defining and essential characteristic of gliomasphere and cancer stem cell phenotype maintenance. (31) (27)

This study was limited by a modest number of samples analyzed. Parallel analysis is a powerful approach to single cell heterogeneity analysis capacities, and 15 samples tested the minimal number of samples needed to quantify these valuable cells for definitive global understanding of this cell subtype. Sample number may have reduced statistical significance of the markers tested and suggested trends, but did not achieve statistically significant prognostic indication, latter particularly a measure to definitively prove these pathways as essential malignancy drivers of these cells (Supplementary Information). (6) (32) PTEN was statistically insignificant in gliomasphere lines and thus proteomic mapping capacities relied on only three markers tested. Despite these limitations, this study did indeed provide insight into the clear utility of quantitative single cell heterogeneity profiling and parallel analysis of these specific pathways contribute to and resolve molecular drivers of the cancer stem-like state and targeting of these cell populations.

## Materials and Methods

### Microfluidic cell array chips

Each cell array chip consists of 24 (3 × 8) chambers, each with dimensions of 8 mm (l) × 1 mm (w) × 120 um (h) and total volume of 960 nL **(Fig. 1B)**. Single cell suspension, culture media, and reagents were introduced and removed at chip ports by electronic, handheld semi-automated pipettor at 6uL/second to protect cells from shear forces and enable flexible reagent and cell control.

### Cell array chip construction

The cell array chip is fabricated by directly attaching a polydimethylsiloxane (PDMS)-based microfluidic component onto an uncoated glass microscope slide. The microfluidic component was fabricated using a soft lithography method. Well-mixed Sylgard 184 PDMS (Corning Inc., A:B = 10:1 ratio) were poured onto a silicon wafer replicate of photolithographically defined microchannel patterns, vacuum degassed and allowed to harden overnight in 80°C oven. The microfluidic component was then peeled off the silicon wafer, edges cut with razor tool (Stanley, Inc.) and holes punched with pipette tip size-matched diameters at the ends of the microchannels. Direct attachment to uncoated glass slide was accomplished via pretreatment with oxygen plasma of bottom of microfluidic PDMS chip and top of uncoated microscope slide. The assembled chip was then baked in an 80°C vacuum oven for 24 hours. Prior to use, the chips were sterilized by exposure to UV light for 15 minutes.

### Microfluidic chip cell loading and handling

To prepare microfluidic chips for cell loading, matrigel at 1:20 (BD Biosciences, Inc.) was used as a cell capture reagent and loaded into chambers for 12 hours at 8°C then washed with PBS. Although precise cell densities at time of loading were dependent on individual sample gliomasphere growth characteristics, cells were dissociated using TrypLE (Invitrogen), spun down at 1200 RPM for 1 min and pelleted, then resuspended at a density of 50 to 500 cells/μL for a 100-1000 μL aliquot containing single cells with media in a 1.5mL tube (Eppendorf, Inc.). The tubes were triturated with Matrix pipettor and 2uL of cell suspension was loaded per matrigel pretreated microfluidic chamber. Three chambers were loaded per gliomasphere sample. Chips were then spun at 1000 RPM for 1 minute to assure all cells would fall into the same Z-plane for imaging. Chips were then placed in a 10-cm Petri dish with 1 mL double-distilled water (for hydration) and incubated in a 5% CO_2_, 37°C incubator for 10 minutes prior to on-chip quantitative immunocytometry.

### Gliomasphere models

Collection of patient tumor tissue for the derivation of gliomaspheres was approved by the Institutional Review Board of UCLA. Briefly, tumors were washed, minced with a scalpel blade, digested with TrypLE (Invitrogen) for 5 minutes and spun down at 1200 RPM for 5 minutes. TrypLE was removed and tumor pieces were resuspended in chilled DMEM-F12 (Invitrogen), dissociated with at least 2 glass pasteur pipets (Fisher Brand) fire polished to successively smaller bores and put through a 70μm and 40um cell strainers (BD Biosciences). A Percol (GE Health Sciences) protocol was employed to remove red blood cells. (6)

Cells were seeded at a density of 100,000 cells/mL in a stem cell growth and enrichment medium consisting of DMEM/F12 medium supplemented with 1:50 B27 (Invitrogen), 20 ng/ml bFGF (Peprotech), 50 ng/mL EGF (Peprotech), 1:100 penicillin/streptomycin (Invitrogen), 1:100 Glutamax (Invitrogen) and 5 ug/ml heparin (Sigma-Aldrich). Heparin, bFGF and EGF were supplemented weekly and Glutamax bi-weekly. Media was changed upon passaging or when media became acidic. Passaging was done according to visual observation when the majority of neurosphere aggregates were observed to merge into larger aggregates. Spheres were passaged into fresh media following either enzymatic dissociation with TrypLE and glass pipet dissociation or chopping using an McElwin automated tissue chopper (Geneq. Inc.). (1)(6)

### Quantitative microscopic imaging

Optimization protocols were developed for assessment of phosphorylation of ribosomal protein S6 for EGFR (Hylite 750nm) and PTEN (555nm) and activated downstream phosphorylation of AKT (Alexa 647nm) and TORC1 (via readout of activated phosphorylated S6 (Alexa 488nm)) (See **Supplemental Methods**). Microfluidic cell array chips facilitated cell and reagent control and improved signal-to-noise ratio. Each chip accommodates 3 chambers/sample and up to 8 samples on each microfluidic chip **(Fig. 1B).** Details on optimization procedures and imaging are available in **Supplementary Methods**. Individual images were taken for the 4 fluorophore-labeled antibodies (488nm, PE, 647nm and 750nm). (20)

Chips containing fixed immunolabelled cells were mounted onto a Nikon TE2000S inverted fluorescent microscope with a CCD camera (Photometrics, Inc.) and X-cite light source (Lumen Dynamics Group). The size of each channel had design specifications for edges to align outside the imaging area. Each channel had a length permitting 8 imageable frames, and all frames were used for image analysis. Imaging parameters were optimized and controlled for to assure all data could be directly comparable. The light source bulb was evaluated in between each sample imaged for operational fidelity. Quantitative imaging was obtained by measuring the fluorescence intensity for each cell area using MetaMorph (Molecular Devices, Version 7.5.6.0). Description of the MetaMorph software module are available in **Supplementary Methods.**

### Bioinformatic self organizing maps (SOMs) and clustering derivation

Self-organizing maps (SOMs) were generated in R (**Supplemental Methods).** (20,33) A SOM grid consists of a set of units each characterized by a codebook vector consisting of the four values (EGFR, PTEN, pAKTAKT and pS6). Input measurements were unit normalized. The codebook vectors are then randomly initialized based on the input data and a training process involves repeated presentation of the training data to the map. Each presented datapoint is assigned to a most similar “winning” grid and the codebook vector of the winning grid is updated using a weighted average, where the weight is the learning rate α. Three SOMs are trained for each data set, and the resulting maps examined for qualitative consistency. Testing various SOM grid sizes identified a 7 × 7 grid as smallest size to capture differences between gliomasphere samples. **(Fig. 2D).**

Hierarchical clustering of Neighborhood Frequency Vectors (NFVs) of SOMs **(Fig. 2E**, **3A)** with waterfall plots displaying differing average intensities values for each cluster were generated **(Fig. 2B, 2F)**. Further analytical details can be found in the **Supplemental Methods**.

### TCGA Microarray Analysis

The Verhaak et al. classification of The Cancer Genome Atlas Glioblastoma database was used to inform TCGA analysis. (10) The unified gene expression dataset is the combined expression data from three platforms, Affymetrix HuEx array, Affymetrix U133A array and Agilent 244K array into a single expression pattern that was used for the original classification of the TCGA dataset into four categories. The unified gene expression data was combined with tumor and gliomasphere data which was obtained on the Affymetrix U133 plus 2.0 array and normalized with the using the R package limma. (34,35) Batch effects were then adjusted using ComBat (36) on the normalized data. ClaNC, the LDA based centroid classification algorithm used by Verhaak et al. to create the classifications was then applied to determine a 3-class centroid-based classifier using only the data from Mesenchymal, Proneural or Classical TCGA samples. (37) The original dataset has been reported on previously, consisting of 56 Mesenchymal samples, 53 Proneural and 38 Classical samples consisting of 147 total samples excluding the 26 Neural samples were used in building the classifier. (11) This classifier was then used to assign a TCGA category (Mesenchymal, Proneural or Classical) to each gliomasphere sample. Because of the lack of gene name overlap from the Affymetrix U133A array used by TCGA and the Affymetrix U133 plus 2.0 microarray used for our classifications, only 789 of the original 840 genes were used to classify the samples.

### Quantitative neurosphere analysis

Neurosphere size and number measurements were obtained with an Acumen eX3 plate reader in the UCLA California NanoSystems Institute (CNSI) Molecular Shared Screening Resource core facility. For this, cells were fixed with 1:1 mixture of 4% paraformaldehyde and 100% methanol, 50 μL/well. After at least 4 hrs post-fixation the DNA binding Syto-9 dye was added (1:1000 dilution in PBS, 10 μL/well). The parameters from data output for each identified object included peak and total intensities (FLU), diameter (expressed as width and depth in μm), and spherical volume (μm3). Thresholds for false, i.e. non-neurosphere objects (e.g., cell clumps, single cells, DNA remnants, debris, etc), were defined by: 1) objects with diameter <35 um, 2) peak intensity <100 FLU or 3) width/depth ratio >4. After setting thresholds, means and standard error for sphere numbers were calculated based on number of objects from an average of 10 wells and mean spherical volume per condition to estimate neurosphere size. (38)

### Erlotinib LC_50_

Assessment of LC_50_ via quantitative measurements of sphere size and sphere number was deployed (**Fig. 3B)**. Gliomaspheres were dissociated and resuspended in neurosphere media and plated into 96 well microplates at a 50 cell/well density for each sample (10 wells/condition). Experimental parameters included DMSO treated control wells and 5 conditions treated with serial dilutions of Erlotinib (LC Laboratories) to a final volume 100 μL/well. Final DMSO concentration was equalized to match with DMSO% in highest concentration for each compound. Plates were incubated and monitored for formation of 10 neurospheres/well, occurring at approximately 16 days post-incubation.

### Sphere initiation efficiency assays

Limiting dilution assays were performed by single cell dissociation and resuspension in neurosphere media and plated into 96 well microplates. A measure of sphere forming efficiency was achieved by seeding incrementally increasing numbers of cells at intervals 50 cells up to 800 cells/well and assessing the number of cells required to achieve ten gliomasphere spheres per well. Plates were incubated and monitored for sphere formation over a 16 day incubation period. The minimal cell density to achieve 10 gliomasphere neurospheres per well is reported **(Fig. 3C)**.

### Patient analyses

This study was overseen and approved by the UCLA Institutional Review Board and HIPAA compliant. Patient demographics, treatment and outcome data are available in **Supplemental Table 1**. Eligible patients consisted of full treatment for glioblastoma, including surgery, chemotherapy and/or radiation therapy and resected tissues capable of renewable neurosphere formation and maintained for at least three passages. Outcome and survival curves with their corresponding hazard ratios are available in **Supplementary Methods**.

## Abbreviations

TBD: 

## Acknowledgements

All experiments with human cells were conducted under protocols approved and overseen by the Institutional Review Board of the UCLA Office of Protection of Research Subjects. This study was supported by the National Natural Science Foundation of China (No. 21505013), the Natural Science Foundation of Liaoning Province, China (No. 2015020660), and the Dalian Science and Technology Innovation Funds (No. 2018J13SN087).

## References

1. Batlle E, Clevers H. Cancer stem cells revisited. Nature Medicine 2017;23:1124

2. Nimmakayala RK, Batra SK, Ponnusamy MP. Unraveling the journey of cancer stem cells from origin to metastasis. Biochimica Et Biophysica Acta-Reviews on Cancer 2019;1871:50–63

3. Delgado-López PD, Corrales-García EM. Survival in glioblastoma: a review on the impact of treatment modalities. Clinical and Translational Oncology 2016;18:1062–71

4. Phillips HS, Kharbanda S, Chen R, Forrest WF, Soriano RH, Wu TD, et al. Molecular subclasses of high-grade glioma predict prognosis, delineate a pattern of disease progression, and resemble stages in neurogenesis. Cancer Cell 2006;9:157–73

5. Chen J, Li Y, Yu T-S, McKay RM, Burns DK, Kernie SG, et al. A restricted cell population propagates glioblastoma growth after chemotherapy. Nature 2012;488:522–6

6. Laks DR, Masterman-Smith M, Visnyei K, Angenieux B, Orozco NM, Foran I, et al. Neurosphere formation is an independent predictor of clinical outcome in malignant glioma. Stem Cells 2009;27:980–7

7. Brennan Cameron W, Verhaak Roel GW, McKenna A, Campos B, Noushmehr H, Salama Sofie R, et al. The Somatic Genomic Landscape of Glioblastoma. Cell 2013;155:462–77

8. Dey-Guha I, Wolfer A, Yeh AC, G. Albeck J, Darp R, Leon E, et al. Asymmetric cancer cell division regulated by AKT. Proceedings of the National Academy of Sciences 2011;108:12845

9. Srivas R, Shen JP, Yang CC, Sun SM, Li J, Gross AM, et al. A Network of Conserved Synthetic Lethal Interactions for Exploration of Precision Cancer Therapy. Molecular cell 2016;63:514–25

10. Verhaak RGW, Hoadley KA, Purdom E, Wang V, Qi Y, Wilkerson MD, et al. Integrated genomic analysis identifies clinically relevant subtypes of glioblastoma characterized by abnormalities in PDGFRA, IDH1, EGFR, and NF1. Cancer cell 2010;17:98–110

11. Laks DR, Crisman TJ, Shih MYS, Mottahedeh J, Gao F, Sperry J, et al. Large-scale assessment of the gliomasphere model system. Neuro-oncology 2016;18:1367–78

12. Jiang H, Gao HP, Kong J, Song B, Wang P, Shi BZ, et al. Selective Targeting of Glioblastoma with EGFRvIII/EGFR Bitargeted Chimeric Antigen Receptor T Cell. Cancer Immunology Research 2018;6:1314–26

13. Gedeon PC, Schaller TH, Chitneni SK, Choi BD, Kuan CT, Suryadevara CM, et al. A Rationally Designed Fully Human EGFRvIII:CD3-Targeted Bispecific Antibody Redirects Human T Cells to Treat Patient-derived Intracerebral Malignant Glioma. Clin Cancer Res 2018;24:3611–31

14. Bhat KPL, Balasubramaniyan V, Vaillant B, Ezhilarasan R, Hummelink K, Hollingsworth F, et al. Mesenchymal differentiation mediated by NF-κB promotes radiation resistance in glioblastoma. Cancer cell 2013;24:331–46

15. Cusulin C, Chesnelong C, Bose P, Bilenky M, Kopciuk K, Chan JA, et al. Precursor States of Brain Tumor Initiating Cell Lines Are Predictive of Survival in Xenografts and Associated with Glioblastoma Subtypes. Stem cell reports 2015;5:1–9

16. Furnari FB, Cloughesy TF, Cavenee WK, Mischel PS. Heterogeneity of epidermal growth factor receptor signalling networks in glioblastoma. Nat Rev Cancer 2015;15:302

17. Verhaak RG, Hoadley KA, Purdom E, Wang V, Qi Y, Wilkerson MD, et al. Integrated genomic analysis identifies clinically relevant subtypes of glioblastoma characterized by abnormalities in PDGFRA, IDH1, EGFR, and NF1. Cancer Cell 2010;17:98–110

18. Tome-Garcia J, Erfani P, Nudelman G, Tsankov AM, Katsyv I, Tejero R, et al. Analysis of chromatin accessibility uncovers TEAD1 as a regulator of migration in human glioblastoma. Nat Comm 2018;9:13

19. Capuani F, Conte A, Argenzio E, Marchetti L, Priami C, Polo S, et al. Quantitative analysis reveals how EGFR activation and downregulation are coupled in normal but not in cancer cells. Nat Comm 2015;6:7999

20. Sun J, Masterman-Smith MD, Graham NA, Jiao J, Mottahedeh J, Laks DR, et al. A microfluidic platform for systems pathology: multiparameter single-cell signaling measurements of clinical brain tumor specimens. Cancer Res 2010;70:6128–38

21. Leu S, von Felten S, Frank S, Vassella E, Vajtai I, Taylor E, et al. IDH/MGMT-driven molecular classification of low-grade glioma is a strong predictor for long-term survival. Neuro-Oncology 2013;15:469–79

22. Farhy C, Hariharan S, Ylanko J, Orozco L, Zeng F-Y, Pass I, et al. Improving drug discovery using image-based multiparametric analysis of the epigenetic landscape. eLife 2019;8:e49683

23. Lin D, Li P, Feng J, Lin Z, Chen X, Yang N, et al. Screening Therapeutic Agents Specific to Breast Cancer Stem Cells Using a Microfluidic Single-Cell Clone-Forming Inhibition Assay. Small;0:1901001

24. Filbin MG, Tirosh I, Hovestadt V, Shaw ML, Escalante LE, Mathewson ND, et al. Developmental and oncogenic programs in H3K27M gliomas dissected by single-cell RNA-seq. Science 2018;360:331–5

25. Hu AX, Adams JJ, Vora P, Qazi M, Singh SK, Moffat J, et al. EPH Profiling of BTIC Populations in Glioblastoma Multiforme Using CyTOF. In: Singh SK, Venugopal C, editors. Brain Tumor Stem Cells: Methods and Protocols. New York, NY: Springer New York; 2019. p 155–68.

26. Lan X, Jörg DJ, Cavalli FMG, Richards LM, Nguyen LV, Vanner RJ, et al. Fate mapping of human glioblastoma reveals an invariant stem cell hierarchy. Nature 2017;549:227

27. Kelly JP, Stechishin O, Chojnacki A, Lun X, Sun B, Senger DL, et al. Proliferation of human glioblastoma stem cells occurs independently of exogenous mitogens. Stem Cells 2009;27:1722–33

28. Wei W, Shin YS, Xue M, Matsutani T, Masui K, Yang H, et al. Single-Cell Phosphoproteomics Resolves Adaptive Signaling Dynamics and Informs Targeted Combination Therapy in Glioblastoma. Cancer cell 2016;29:563–73

29. Luchman HA, Stechishin ODM, Nguyen SA, Lun XQ, Cairncross JG, Weiss S. Dual mTORC1/2 blockade inhibits glioblastoma brain tumor initiating cells and synergizes with temozolomide to increase orthotopic xenograft survival. Clinical Cancer Research 2014;20:5756–67

30. Euskirchen P, Radke J, Schmidt MS, Schulze Heuling E, Kadikowski E, Maricos M, et al. Cellular heterogeneity contributes to subtype-specific expression of ZEB1 in human glioblastoma. PloS one 2017;12:e0185376–e

31. Han Y-P, Enomoto A, Shiraki Y, Wang S-Q, Wang X, Toyokuni S, et al. Significance of low mTORC1 activity in defining the characteristics of brain tumor stem cells. Neuro-oncology 2017;19:636–47

32. Panosyan EH, Laks DR, Masterman-Smith M, Mottahedeh J, Yong WH, Cloughesy TF, et al. Clinical outcome in pediatric glial and embryonal brain tumors correlates with in vitro multi-passageable neurosphere formation. Pediatr Blood Cancer 2010;55:644–51

33. Wehrens R, Buydens LMC. Self- and super-organizing maps in R: The Kohonen package. J Stat Soft 2007:1–19

34. Smyth GK. limma: Linear Models for Microarray Data. In: Gentleman R, Carey VJ, Huber W, Irizarry RA, Dudoit S, editors. Bioinformatics and Computational Biology Solutions Using R and Bioconductor. New York, NY: Springer New York; 2005. p 397–420.

35. Rc T. 2013 R, A language and environment for statistical computing. <http://www.R-project.org/>.

36. Johnson WE, Li C, Rabinovic A. Adjusting batch effects in microarray expression data using empirical Bayes methods. Biostatistics 2007;8:118–27

37. Dabney AR. ClaNC: point-and-click software for classifying microarrays to nearest centroids. Bioinformatics 2006;22:122–3

38. Visnyei K, Onodera H, Damoiseaux R, Saigusa K, Petrosyan S, De Vries D, et al. A molecular screening approach to identify and characterize inhibitors of glioblastoma stem cells. Mol Cancer Ther 2011;10:1818–28

