## Supplementary Information for "Quantitative single cell heterogeneity profiling of patient derived tumor initiating gliomaspheres reveals unique signatures of drug response and malignancy"

**Supplemental Information**

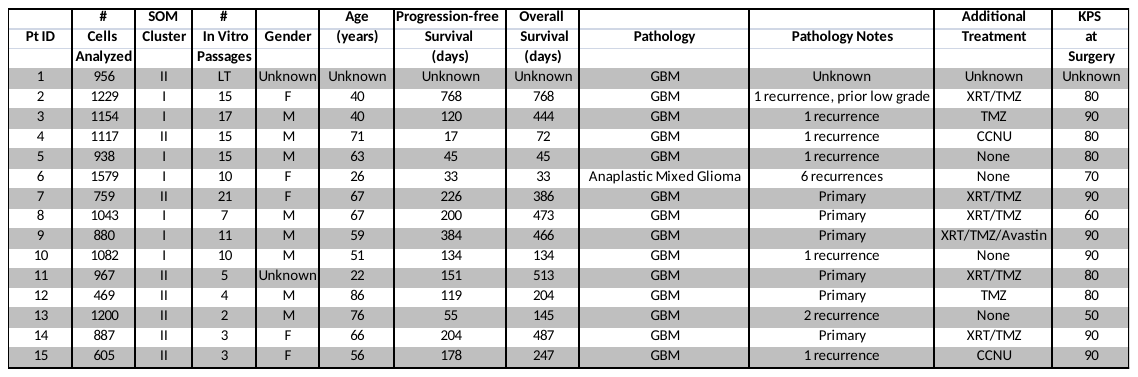

**Supplemental Table 1. Patient data.**

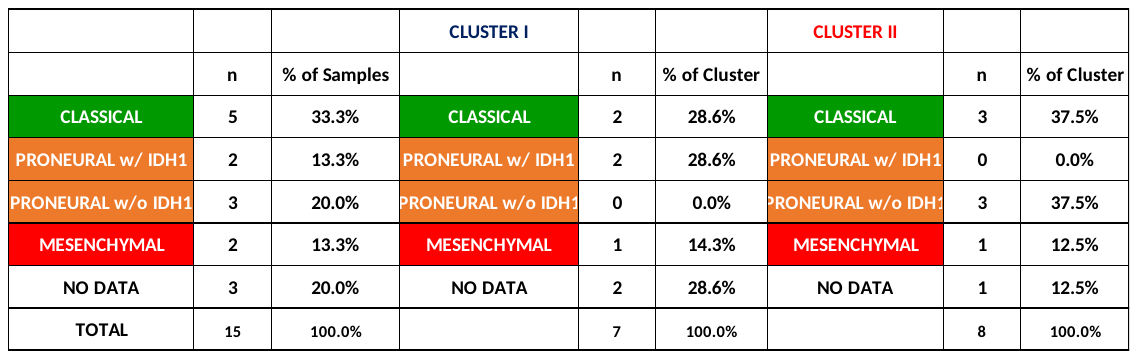

**Supplemental Table 2. Distribution of TCGA subtypes.**

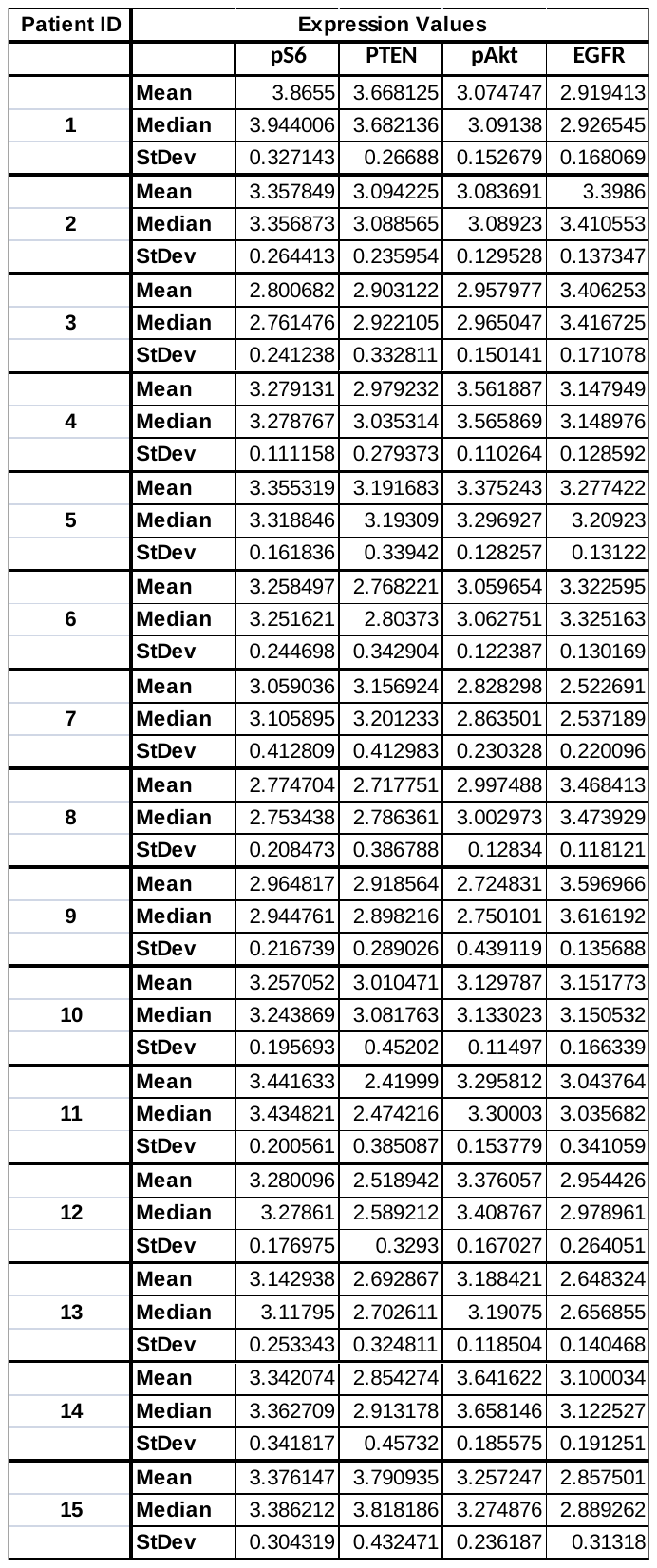

**Supplemental Table 3. Mean, Median, Standard Deviation (StDev) expression of each biomarker from each patient.**

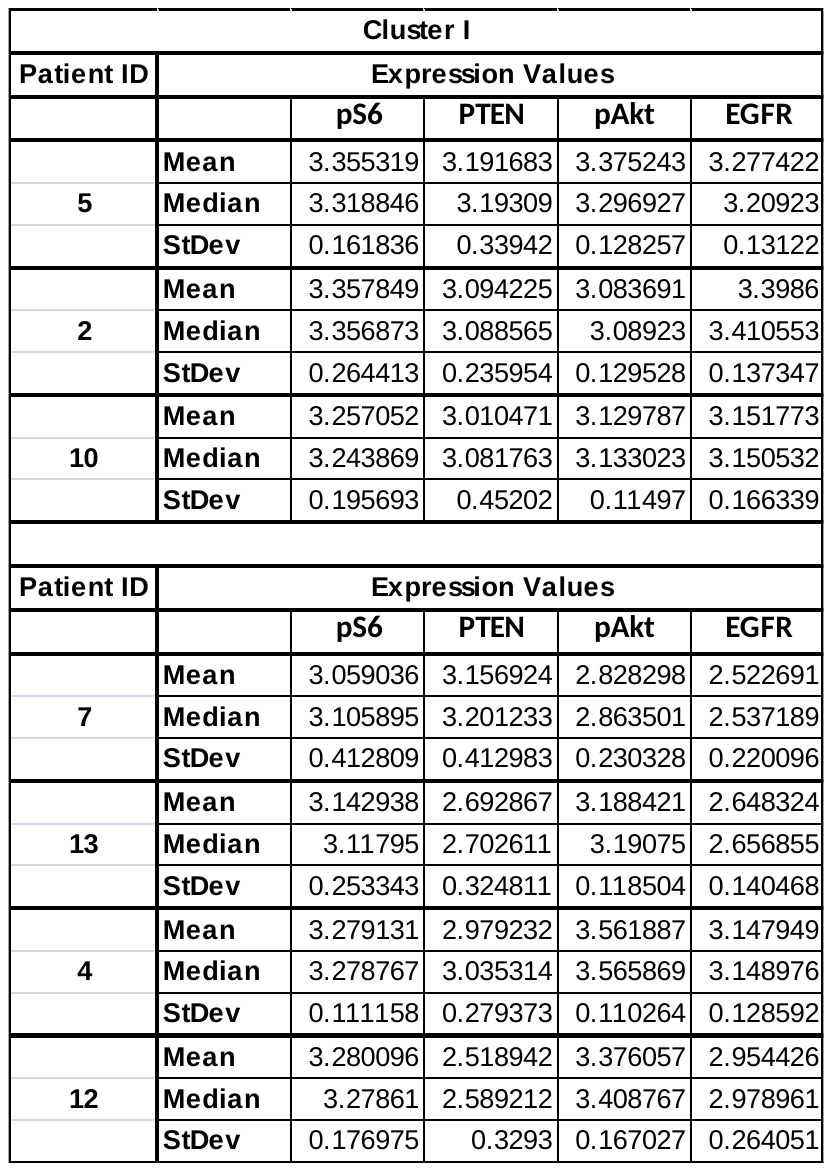

**Supplemental Table 4a. Mean, median, standard deviation (StDev) of expression values of all drug-treated patient samples.**

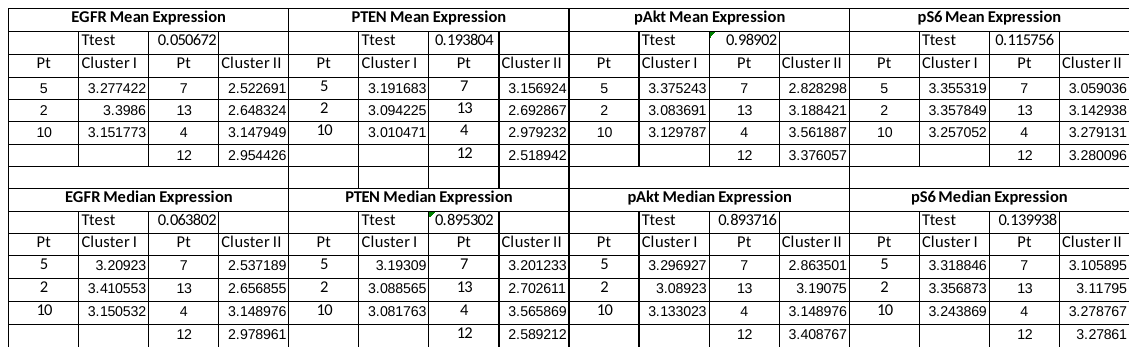

**Supplementary Table 4b. Average expression of erlotinib drug-treated samples.** Student’s T-test on expression of all drug-treated patient samples. Borderline significant differences were found for EGFR expression.

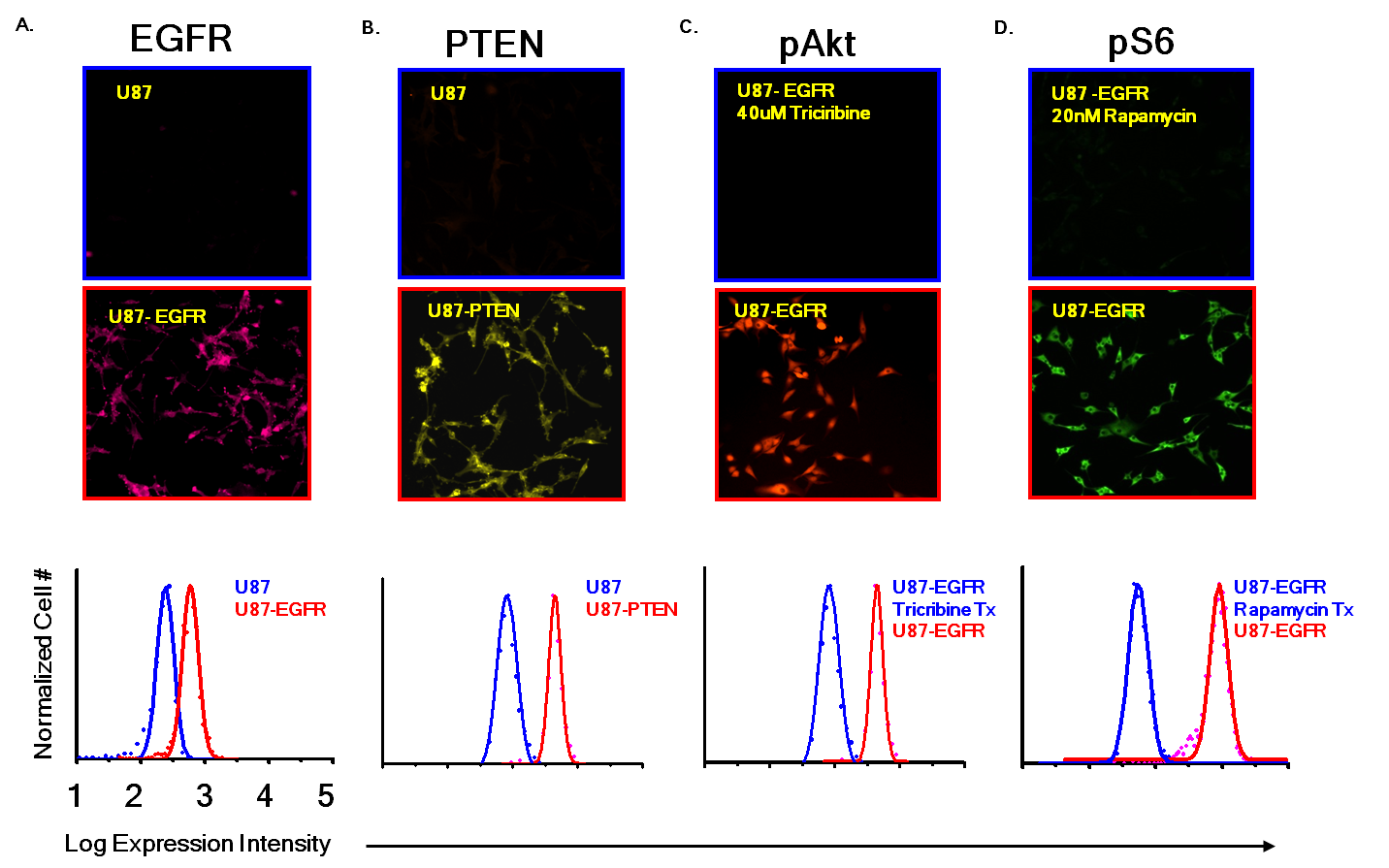

**Supplemental Figure 1. Optimization EGFR, PTEN, pAkt, and pS6 signalling assay.** Immunofluorescent images of immunocytochemistry (top) negative controls (middle) positive controls and (bottom) quantitative histograms of the dynamic ranges of single cell measurements. **Figure 1a-b**. The U87 glioblastoma cell line was used as a negative control for EGFR and PTEN analysis and the U87-EGFR overexpression variant was used as the positive control for EGFR and U87-PTEN overexpression variant was used as the positive control for PTEN. **Figure 1c-d.**

U87-EGFR with inhibitor-treated lines used for pAkt and pS6 analysis. (Bottom) X-axes, logarithmic transformed expression intensity. Y-axes, normalized cell numbers. Blue, negative control. Red, positive control.

*Optimization of immunoassaying parameters*

The U87 glioblastoma cell line and genetically modified overexpression variants identified the appropriate antibody concentrations to identify the maximum range in average signal intensity between positive and negative controls. Optimization of the immunoassay was accomplished by quantifying the expression of each biomarker in control cell lines.

Cell lines were maintained in DMEM-F12 (Invitrogen) containing 10% fetal bovine serum (HiClone) and 1% penicillin/streptomycin (Invitrogen). For EGFR, the positive control was a U87 glioblastoma line genetically modified to overexpress EGFR and standard U87 cells served as negative control **(Supplementary Figure 1a)**. For PTEN, the positive control was a U87 glioblastoma line genetically modified to overexpress PTEN with standard U87 cells as the negative control **(Supplementary Figure 1b)**. For activation of Akt and S6, U87 cells overexpressing EGFR were used as positive controls and the pAkt inhibitor tricribine (LC Laboratories) **(Supplementary Figure 1c)** and the mTOR inhibitor rapamycin (Sigma) **(Supplementary Figure 1d)** were used in conjunction with the U87-EGFR expression variant as the negative control for optimization of pAkt and pS6, respectively. Cells were treated with 40 μmol/L triciribine for 48 hours, 20 nmol/L rapamycin for 48 hours.

*On-chip immunocytochemistry*. To prepare cells for loading and immunoassaying in microfluidic channels, cells were dissociated using TrypLE (Invitrogen), pelleted, and resuspended at a density of 250 to 500 cells/μL. Two microliters of the U87 cell suspension were loaded into each microfluidic channel. Chips were then placed in a 10-cm Petri dish with 1 mL double-distilled water (for hydration) and incubated in a 5% CO_2_, 37°C incubator for 16 hours before drug treatment and on-chip ICC. On-chip ICC involved cell fixation (4% paraformaldehyde for 15 min at RT), washing with PBS, cell permeabilization (0.3% Triton X-100 for 15 minutes at RT), washing, blocking (10% normal goat serum, 3% BSA and 0.1% N-doceyl-B-maltodextrose) for 12 hours at 4°C, immunolabeling for 12 hours at 4°C followed by washing and DAPI staining prior to imaging. Multiparameter immunolabeling was achieved with an optimized mixture of fluorophore-conjugated antibodies prepared by mixing the empirically derived concentrations (Supplementary Figure 1) of 2.754 μg/mL anti-EGFR (BD Pharmingen) labeled with LiCor/HiLyte Fluor 750 labeling kit (Dojindo Molecular Technologies, Inc.), 0.625 μg/mL PE-conjugated anti-PTEN (BD Biosciences), 1.250 μg/mL Alexa Fluor 647-conjugated anti-pS473-AkT (Cell Signaling Technology) and 3.800 μg/mL Alexa Fluor 488-conjugated anti-pS235/S236-S6 (Cell Signaling Technology).

*Imaging*. Exposure times were optimized and identified: 10 sec for HiLyte Fluor 750 (EGFR), 1 second for PE (PTEN), 2 seconds for Alexa Fluor 647 (pAkt) and 0.5 seconds for Alexa Fluor 488 (pS6). DAPI staining allowed the area of nuclei to be measured and counted.

*Quantitative image analysis*. MetaMorph (Molecular Devices, Version 7.5.6.0) *Multi-Wavelength Cell Scoring* module allowed image analysis for expression intensity scoring and quantification of cells. After setting minimum and maximum cell size thresholds, Metamorph identifies “cell regions,” which were inspected by the user to verify that all cells (but not cell debris) were accurately identified by the Metamorph algorithm. Background subtraction for each frame was performed by assessing the average intensity values in areas with no cells and subtracting this intensity value from each cell's score for each fluorescent signal. Integrated fluorescent intensity values were then divided by the cell area to achieve cell-spread surface-area average intensities, and these values were logarithmically transformed to give Gaussian-like distributions. These log-transformed values were used for subsequent analysis.

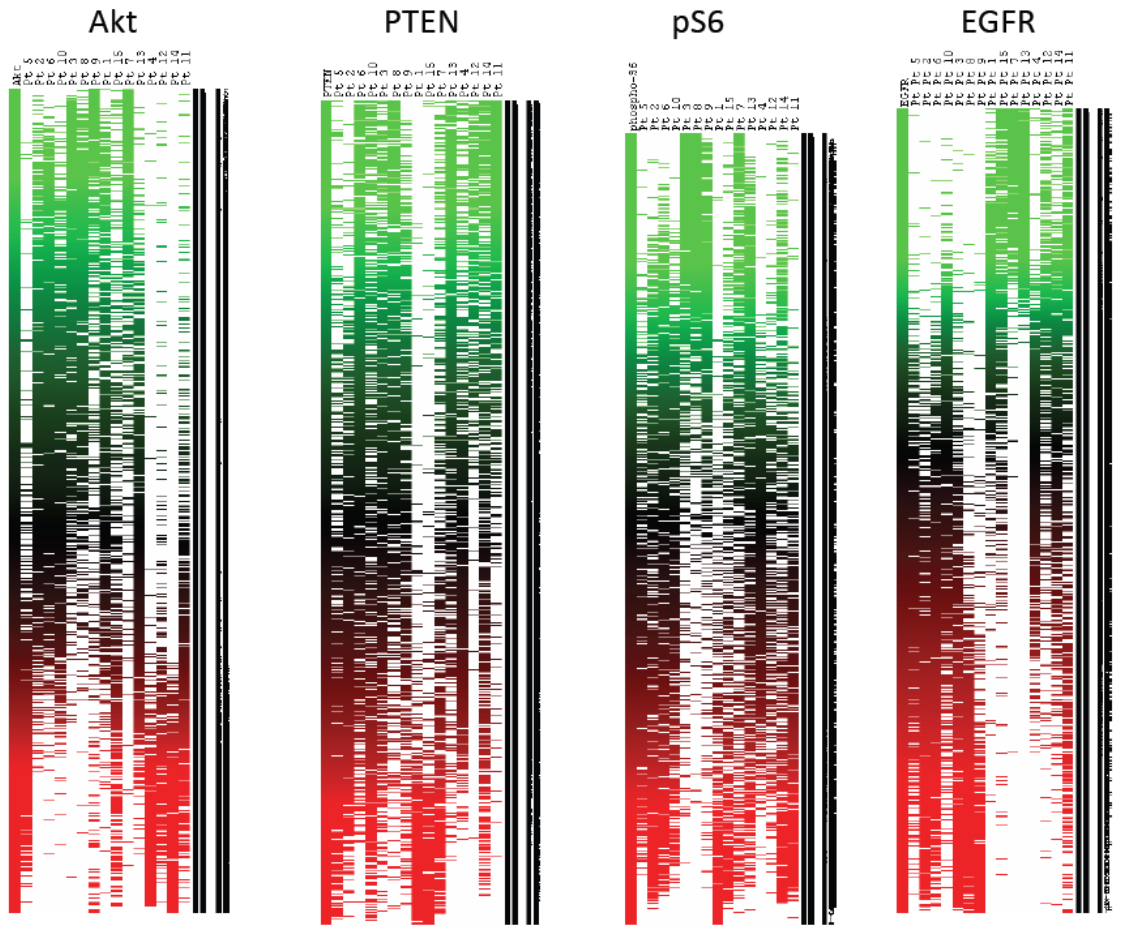

**Supplemental Figure 2. Single cell expression heatmps for Akt, PTEN, pS6, EGFR. Red, high expression. Green, low expression.**

**Bioinformatic analysis**

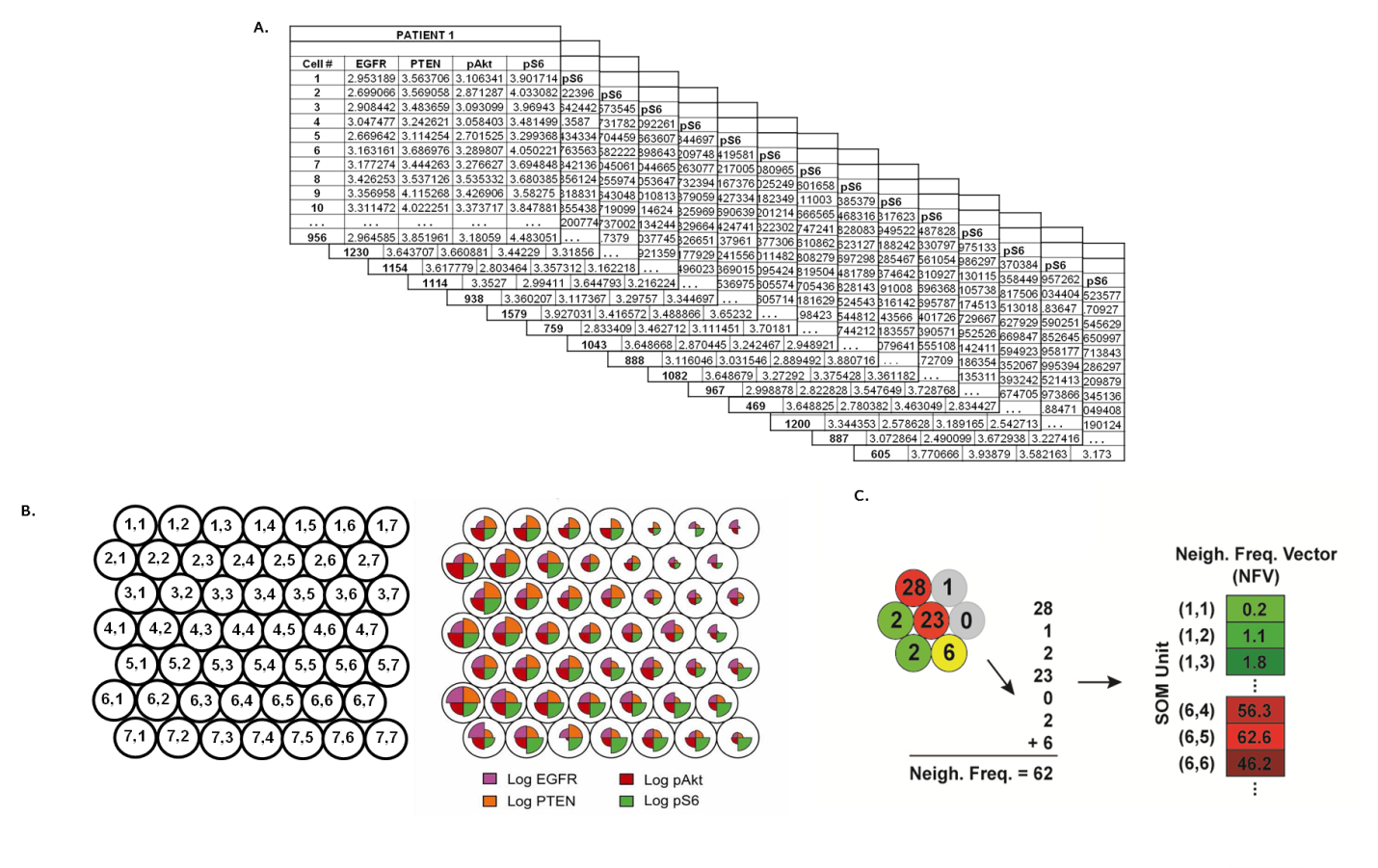

**Supplemental Figure 3. Bioinformatic Analysis.**

A. Excel spreadsheets (truncated for visualization purposes) displaying 4 individual expression values for each biomarker in each cell in 15 patient lines. Columns, EGFR, PTEN, pAkt, pS6. Rows, individual cells. B. Self-organizing map (SOM). (Left) 7x7 grid allows grouping of individual cells into 49 molecular unique profiles. (Right) Pie chart representation of grid characteristics. C. Quantitatively comparison of SOM mappings of line subject to neighborhood frequency vectoring (NFV) is calculated. (Left) NFV process consists of adding percentage of individual SOM unit and neighboring units. (Right) example of NFV-derived data used in hierarchical clustering. The mean values for all four stains plotted against one another.

For SOM training, the input data set is a compendium of many measurements that represent the global measurement space. Here, the input training data set consisted of ~15,000 single-cell 4-parameter measurements from 15 specimens (469-1579 cells per sample). After training, the individual samples from the compendium were then be mapped to the SOM to indicate which portion of the parameter space is occupied by an individual sample. This mapping procedure involves assessment of the best-matching SOM unit for each single-cell 4-parameter measurement in the data set, followed by assignment of that single-cell measurement to the best-matching unit. Running R Version 2.8.1 on a desktop PC with a 2.66 GHz processor, SOM training and mapping of the patient sample data set required less than 1 minute of computational time. To normalize for different number of cells analyzed across data sets, we then convert the number of cells in each unit to a frequency. Thus, regions of a SOM mapping that have a large frequency of cells indicate that many cells within that data set closely resemble the 4-value codebook vector that characterizes that unit.

*SOM Construction*

Using R Version 2.8.1, the Kohonen package for creation and analysis of self-organizing maps was downloaded, installed and loaded.^1, 2^ Quantitative ICC measurements from each individual patient were aggregated into a global data set of 54,884 4-parameter measurements and mean normalized. Using the *som* command in R, a 7x7 hexagonally-packed SOM was trained using the mean-normalized data set with 100 iterations and a learning rate (〈) of 0.05. Because each training process involves random initialization of the codebook vectors and thus distinct map topologies, three SOMs were trained for each data set, and the resulting maps were examined for qualitative consistency. Running R Version 2.8.1 on a desktop PC with a 2.66 GHz processor, SOM training and mapping of the patient sample data set required less than 1 minute of computational time. The resulting SOMs were checked for: 1) convergence (as measured by the average distance of an object with the closest codebook vector unit) and 2) quality (the mean similarity of objects mapped to a unit to the codebook vector of that unit).

*Mapping of patient samples to the SOM*

For a given SOM, the *map* command in R was used to map the measurements for each individual patient to the SOM. Using the *hist* command, the number of cells mapped to each SOM unit was calculated and then converted to a frequency. The log2 frequencies were plotted on the SOM grid using a rainbow color scale. This mapping process was repeated for each of the three trained SOMs, and individual patient sample mappings were examined for qualitative consistency.

**Supplemental Figure 4a. SOM Generation.** Fifteen samples mapped to SOMs and clustering with 100-NFVs (density of cell + 100% of cell’s neighbors)

**Frequency (%)**

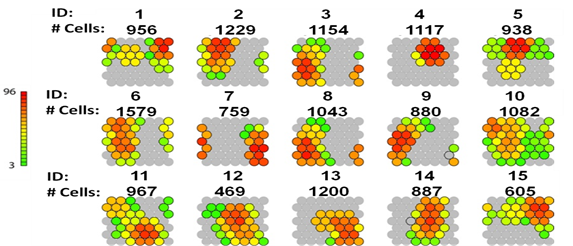

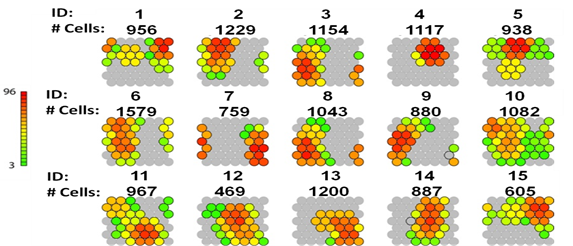

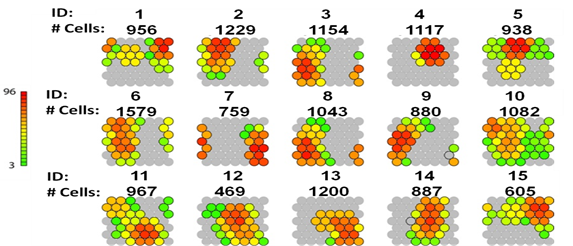

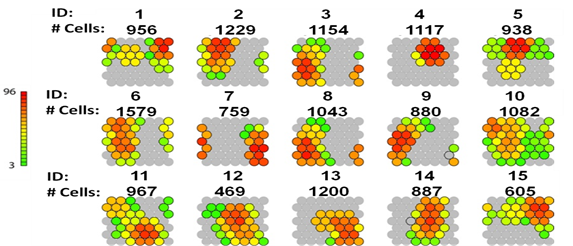

**1	2	3	4	5**

**956	1229	1154	1117	938**

**6	7	8	9	10**

**1579	759	1043	880	1082**

**11	12	13	14	15**

**1579	469	1200	887	605**

**Patient ID:**

**# Cells:**

**Patient ID:**

**# Cells:**

**Patient ID:**

**# Cells:**

NFVs are constructed with same methods as published previously. (1) – the density of each cell plus the sum of the densities of that cell’s neighbors.

*Clustering of SOM mappings*

For each patient sample mapping, the Neighborhood Frequency Vector was calculated. These NFVs were subjected to unsupervised hierarchical clustering using the average-linkage method based on the Pearson correlation using Cluster software. Data were visualized using Java Treeview 1.1.3.^3^

**Kaplan Meier Curves**

Kaplan-Meier distributions for patient time-to-progression (TTP) and time-to-survival (TTS) are plotted in **Supplemental Figure 4b**. A Cox proportional hazards model as implemented in Stata 8.0 (StataCorp) was employed to relate cluster variables. TTP is the duration of progression free survival from the date of surgery until recurrence, death, or, if progression free, until the last follow up date. TTS is the duration of life from the date of surgery. Progression and survival data were unavailable from one patient (Patient 1). Bioinformatic outliers were removed from progression and survival analyses by the criterion of failing to cluster with at least two samples (Patients 5 and 11).

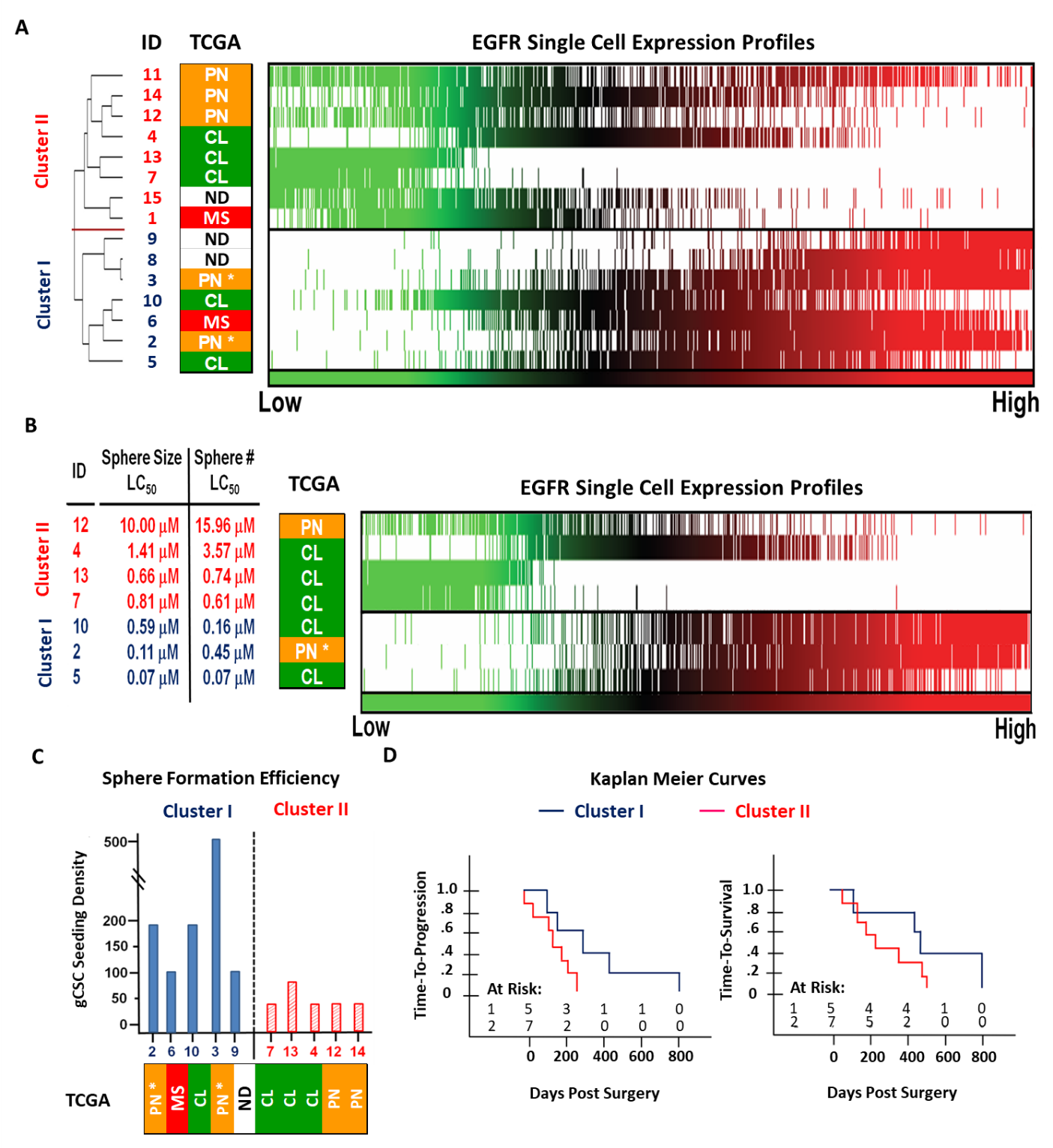

**Supplementary Figure 4b.** Kaplan-Meier plots of patient progression and survival. (Left) Time-to-progression, hazard ratio = 2.93, *p* = 0.14, NS. (Right) Time-to-survival (TTS), survival hazard ratio = 2.51, *p* = 0.16, NS.^1^

**Supplemental Protocols**

*Cell array chip construction*

The cell array chip is fabricated by directly attaching a polydimethylsiloxane (PDMS)-based microfluidic component onto an uncoated glass microscope slide. The microfluidic component was fabricated using a soft lithography method. Well-mixed Sylgard 184 PDMS (Corning Inc., A:B = 10:1 ratio) were poured onto a silicon wafer replicate of photolithographically defined microchannel patterns, vacuum degassed and allowed to harden overnight in 80^o^C oven. The microfluidic component was then peeled off the silicon wafer, edges cut with razor tool (Stanley, Inc.) and holes punched with pipette tip size-matched diameters at the ends of the microchannels. Direct attachment to uncoated glass slide was accomplished via pretreatment with oxygen plasma of bottom of microfluidic PDMS chip and top of uncoated microscope slide. The assembled chip was then baked in an 80^o^C vacuum oven for 24 hours. Prior to use, the chips were sterilized by exposure to UV light for 15 minutes.

*Microfluidic chip-enabled quantitative immunocytochemistry*

To prepare microfluidic chips for cell loading, matrigel at 1:20 (BD Biosciences, Inc.) was used as a cell capture reagent and loaded into chambers for 12 hours at 8^o^C then washed with PBS.

Although precise cell densities at time of loading were dependent on the number and size of spheres in respective samples, cells were dissociated using TrypLE (Invitrogen), spun down at 1200 RPM for 1 min and pelleted, then resuspended at a density of 50 to 500 cells/μL for a 100-1000 μL aliquot containing single cells with media in a 1.5mL tube (Eppendorf, Inc.). The tubes were triturated with Matrix pipettor and 2uL of cell suspension was loaded per matrigel pretreated microfluidic chamber. Three chambers were loaded per gCSC sample. Chips were then spun at 1000 RPM for 1 minute to assure all cells would fall into the same Z-plane for imaging. Chips were then placed in a 10-cm Petri dish with 1 mL double-distilled water (for hydration) and incubated in a 5% CO_2_, 37°C incubator for 10 minutes prior to on-chip quantitative immunocytometry.

On-chip immunocytochemistry involved cell fixation (4% paraformaldehyde for 15 min at RT), washing with PBS, cell permeabilization (0.3% Triton X-100 for 15 minutes at RT), washing, blocking (10% normal goat serum, 3% BSA and 0.1% N-doceyl-B-maltodextrose) for 12 hours at 4°C, immunolabeling for 12 hours at 4°C followed by washing and DAPI staining prior to imaging. Multiparameter immunolabeling was achieved with an optimized mixture of fluorophore-conjugated antibodies prepared by mixing the empirically derived concentrations (Supplementary Figure 1) of 2.754 μg/mL anti-EGFR (BD Pharmingen) labeled with LiCor/HiLyte Fluor 750 labeling kit (Dojindo Molecular Technologies, Inc.), 0.625 μg/mL PE-conjugated anti-PTEN (BD Biosciences), 1.250 μg/mL Alexa Fluor 647-conjugated anti-pS473-AkT (Cell Signaling Technology) and 3.800 μg/mL Alexa Fluor 488-conjugated anti-pS235/S236-S6 (Cell Signaling Technology).

*Microscopic image acquisition and processing*

Chips containing fixed immunolabelled cells was mounted onto a Nikon TE2000S inverted fluorescent microscope with a CCD camera (Photometrics, Inc.) and X-cite light source (Lumen Dynamics Group). The size of each channel had design specifications for edges to align outside the imaging area. Each channel had a length permitting 8 imageable frames, and all frames were used for image analysis. Exposure times were 10 sec for HiLyte Fluor 750 (EGFR), 1 second for PE (PTEN), 2 seconds for Alexa Fluor 647 (pAkt) and 0.5 seconds for Alexa Fluor 488 (pS6). DAPI staining allowed the area of nuclei to be measured and counted. Individual images were taken for the 4 fluorophore-labeled antibodies (488nm, PE, 647nm and 750nm). MetaMorph (Molecular Devices, Version 7.5.6.0) was used to quantify fluorescent signals in individual cells. The Multi-Wavelength Cell Scoring module allowed image analysis for expression intensity scoring and quantification of cells. After setting minimum and maximum cell size thresholds, Metamorph identifies “cell regions,” which were inspected by the user to verify that all cells (but not cell debris) were accurately identified by the Metamorph algorithm. The fluorescent intensity for each cell area was quantified. Background subtraction for each frame was performed by assessing the average intensity values in areas with no cells and subtracting this intensity value from each cell's score for each fluorescent signal. Integrated fluorescent intensity values were then divided by the cell area to achieve cell-spread surface-area average intensities, and these values were logarithmically transformed to give Gaussian-like distributions. These log-transformed values were used for subsequent analysis.

*The Cancer Genome Atlas (TCGA) microarray analysis*

The Verhaak et al. classification of The Cancer Genome Atlas Glioblastoma database: The unified gene expression dataset is the combined expression data from all three platforms, Affymetrix HuEx array, Affymetrix U133A array and Agilent 244K array into a single expression pattern that was used for the original classification of the TCGA dataset into four categories by Verhaak et al. ^4^ The unified gene expression data was combined with tumor and gCSC data which was obtained on the Affymetrix U133 plus 2.0 array and normalized with the using the R package limma.^5^ Batch effects were then adjusted using ComBat^6^ on the normalized data. ClaNC, the LDA based centroid classification algorithm used by Verhaak et al. to create the classifications was then applied to determine a 3-class centroid-based classifier using only the data from Mesenchymal, Proneural or Classical TCGA samples.^7^ The original dataset consisted of 56 Mesenchymal samples, 53 Proneural and 38 Classical samples consisting of 147 total samples excluding the 26 Neural samples were used in building the classifier. This classifier was then used to assign a TCGA category (Mesenchymal, Proneural or Classical) to each sample within the tumor and gCSC sets. Because of the lack of gene name overlap from the Affymetrix U133A array used by TCGA and the Affymetrix U133 plus 2.0 microarray used for our classifications, only 789 of the original 840 genes were used to classify the samples.

Supplemental Protocols References

1. Team RC. A language and environment for statistical computing. R Foundation for Statistical Computing, Vienna, Austria. <http://www.R-project.org/>*.* 2013

2. Therneau T. A Package for Survival Analysis in S. R package version 2.37-7. <http://CRAN.R-project.org/package=survival>*.* 2014

3. Wehrens R, Buydens LMC. Self- and super-organizing maps in R: The kohonen package. J Stat Softw 2007;21(5):1-19.

4. Verhaak RG, Hoadley KA, Purdom E, et al. Integrated genomic analysis identifies clinically relevant subtypes of glioblastoma characterized by abnormalities in PDGFRA, IDH1, EGFR, and NF1. *Cancer cell.* 2010; 17(1):98-110.

5. Smyth G. *Limma: linear models for microarray data. In: Bioinformatics and Computational Biology Solutions using R and Bioconductor*. New York: Springer; 2005.

6. Johnson WE, Li C, Rabinovic A. Adjusting batch effects in microarray expression data using empirical Bayes methods. *Biostatistics.* 2007; 8(1):118-127.

7. Dabney AR. ClaNC: point-and-click software for classifying microarrays to nearest centroids. *Bioinformatics.* 2006; 22(1):122-123.

8. Visnyei K, Onodera H, Damoiseaux R, Saigusa K, Petrosyan S, De Vries D, et al. A molecular screening approach to identify and characterize inhibitors of glioblastoma Stem Cells. Mol Cancer Ther 2011;10(10):1818-28.
